# SARS-CoV-2 neurotropism-induced anxiety/depression-like behaviors require Microglia activation

**DOI:** 10.1101/2023.10.02.560570

**Authors:** Qian Ge, Shan Zhou, Jose Porras, Panfeng Fu, Ting Wang, Jianyang Du, Kun Li

**Affiliations:** Department of Anatomy and Neurobiology, University of Tennessee Health Science Center, Memphis, TN 38163, USA; Florida Research and Innovation Center, Cleveland Clinic, Port St. Lucie, FL 34987, USA; Neuroscience Institute, University of Tennessee Health Science Center, Memphis, TN, United States; Center for Translational Science, Florida International University, Port St. Lucie, FL, 34987, USA

**Author notes:** Correspondence to: Kun Li, Florida Research and Innovation Center, Cleveland Clinic, Port St. Lucie, FL 34987, USA, Jianyang Du, Department of Anatomy and Neurobiology, University of Tennessee Health Science Center, Memphis, TN 38163, USA. **Authorship note:** Q.G. and S.Z. contributed equally to this work.

**Keywords:** COVID-19, SARS-CoV-2, long COVID, post-acute sequelae of SARS-CoV-2 infection, spike protein, Microglia, Amygdala, neurotropism, anxiety- and depression-like behaviors, synaptic transmission and plasticity

## Abstract

The coronavirus disease 2019 (COVID-19) pandemic, caused by severe acute respiratory syndrome coronavirus 2 (SARS-CoV-2), has been associated with a wide range of “long COVID” neurological symptoms. However, the mechanisms governing SARS-CoV-2 neurotropism and its effects on long-term behavioral changes remain poorly understood. Using a highly virulent mouse-adapted SARS-CoV-2 strain, denoted as SARS2-N501Y_MA30_, we demonstrated that intranasal inoculation of SARS2-N501Y_MA30_ results in viral dissemination to multiple brain regions, including the amygdala and hippocampus. Behavioral assays indicated a marked elevation in anxiety- and depression-like behaviors post infection. A comparative analysis of RNA expression profiles disclosed alterations in the post-infected brains. Additionally, we observed dendritic spine remodeling on neurons within the amygdala after infection. Infection with SARS2-N501Y_MA30_ was associated with microglial activation and a subsequent increase in microglia-dependent neuronal activity in the amygdala. Pharmacological inhibition of microglial activity subsequent to viral spike inoculation mitigates microglia-dependent neuronal hyperactivity. Transcriptomic analysis of infected brains revealed the upregulation of inflammatory and cytokine-related pathways, implicating microglia-driven neuroinflammation in the pathogenesis of neuronal hyperactivity and behavioral abnormality. Overall, these data provide critical insights into the neurological consequences of SARS-CoV-2 infection and underscore microglia as a potential therapeutic target for ameliorating virus-induced neurobehavioral abnormalities.

## INTRODUCTION

As of April, 3rd, 2024, the global pandemic of COVID-19 has resulted in 774,954,393 reported cases and 7,040264 confirmed deaths (WHO COVID-19 Dashboard https://data.who.int/dashboards/covid19/deaths?n=c). The death rate and hospital admissions related to COVID-19 have been dramatically reduced as a result of extensive vaccination rollouts and improved treatments. However, a substantial number of patients (10-20%) are experiencing a persistent or newly developed set of symptoms following the acute phase of the illness. This condition is commonly referred to as post-acute sequelae of SARS-CoV-2 infection (1) also known as “long COVID,” (1). While PASC initially attracted attention for its severe impact on older adults and those with underlying health concerns, it has since been clear that it may also occur in otherwise healthy young people, and it can develop after even a modest initial infection (2, 3). These long-lasting symptoms may persist for weeks or even months, posing challenges for healthcare providers and necessitating further research and support to address the long-term health consequences of the disease.

Among the enduring symptoms of PASC, neurological manifestations stand out prominently. Symptoms include cognitive difficulties, autonomic dysfunction, extreme fatigue, sleep disturbances, and mental health complications such as anxiety and depression (2, 3). The exact cause of these syndromes remains uncertain and is constantly being studied. Possible factors contributing to these symptoms include viral infection of brain cells, immune-mediated phenomena, coincidental events, or a combination of these factors. In vitro studies have provided clear evidence of SARS-CoV-2 infection in human brain organoids or cell cultures (4, 5) Additionally, several autopsy studies have reported the presence of viral RNA or proteins in the brains of patients who died from COVID-19, suggesting the possibility of SARS-CoV-2 neurotropism in the central nervous system (CNS) (5, 6). The neuromechanism of SARS-CoV-2 is not clearly defined, both a systematic mechanism and direct neurotropism have been proposed (7). Recent studies suggest that SARS-CoV-2 triggered the COVID-19 pandemic, and possesses brain neurotropism primarily through binding to the ACE2 on neurons (5), or through tunneling nanotubes (8). In addition to direct invasion of neuronal cells, the activation of microglia and/or astrocytes might contribute to the onset and progression of neurological disorders through abnormal maintenance of homeostasis, leading to altered neuronal activities, and thus is considered critical in defining the neurological damage and neurological outcome of COVID-19 (9).

As the principal innate immune cells of the brain, microglia form the primary focus of research in the field of neuroimmune disease. Microglia are the most dynamic neural cells found to date (10–12). Their profoundly dynamic processes constantly survey the local microenvironment, monitor neuronal activity (13, 14), and respond to infection and injury by releasing pro-inflammatory molecules and phagocytic clearance of apoptotic cells (15). During a homeostatic situation, secretion of molecules and phagocytosis by microglia likewise maintain synaptic transmission and plasticity (16, 17). In contrast, hyperactive microglia can be pathogenic and are associated with symptoms of psychiatric disorders, including anxiety disorders and major depression (18), which have commonly appeared during the COVID-19 pandemic. The activity of microglia can be modulated by neuronal activity (19–21), suggesting the existence of microglia-neuron crosstalk. Despite the high potential relationship between SARS-CoV-2 infection and the neuroimmune system, the mechanisms by which SARS-CoV-2 infection activates pro-inflammatory microglia regulates microglia-neuron interaction and alters neuronal activity have not been adequately studied.

Further research is essential to comprehensively unravel the mechanisms underlying PASC and its neurological manifestations, as this knowledge is critical for developing effective management and treatment strategies for affected patients. Mouse models serve as invaluable tools in conducting such studies. Unfortunately, mice are naturally resistant to ancestral SARS-CoV-2 infection due to the low affinity of the viral spike (S) glycoprotein to mouse ACE2 receptors (mACE2) (22, 23). To overcome this limitation, several strategies have been employed to overexpress human ACE2 (hACE2) in mice. These approaches include the delivery of exogenous hACE2 using a replication-deficient adenovirus (Ad5-hACE2) or Adeno-associated Virus (AAV-hACE2), and the generation of K18-hACE2 transgenic mice (24–26). These models have been used to investigate the acute infection of SARS-CoV-2 as well as the neurological consequences (5). However, it should be noted that these models have drawbacks that limit their application in studying long-term PASC after SARS-CoV-2 infection. The former may be incapable of getting a brain infection, whereas the latter may acquire severe artificial multiorgan infections with a high mortality rate, rendering them poor candidates for long-term post-infection behavioral testing.

A highly virulent mouse-adapted SARS-CoV-2 strain, SARS2-N501Y_MA30_, was successfully isolated through serial passage of a recombinant SARS-CoV-2/spike N501Y virus in BALB/c mice (27). After SARS2-N501Y_MA30_ infection, we observed viral titers, viral genomic RNA (gRNA), and subgenomic RNA (sgRNA) in both the lungs and the brains of C57BL/6 mice, indicating the occurrence of virus infection. Fourteen days after the SARS2-N501Y_MA30_ infection, the mice displayed abnormal anxiety- and depression-like behaviors in multiple behavioral paradigms including the open-field test, elevated plus maze test, tail suspension test, and forced swimming test. Due to the critical role of the amygdala in anxiety and depression-like behaviors in rodents and humans (28), we first examined this region and discovered microglia activation following SARS2-N501Y_MA30_ infection. Furthermore, heightened microglia-dependent neuronal activity in the amygdala was evident upon exposure to the spike protein. Our findings uncover the neurotropic potential of SARS-CoV-2 and its direct link to anxiety- and depression-like behaviors through the activation of microglia-mediated neuroinflammatory pathways. This study sheds valuable insights into the neurological repercussions of SARS-CoV-2 infection and suggests microglia as prospective therapeutic targets for mitigating virus-induced neurobehavioral deficits. Understanding the neurological basis of COVID-19-related neuropsychiatric symptoms is critical for developing effective treatments to reduce the long-term impact of PASC on mental health.

## MATERIALS AND METHODS

### Mice, virus, and cells

Specific pathogen-free 6-9 weeks male and female Balb/c and C57BL/6 mice were purchased from the Jackson Laboratory. All protocols were approved by the Institutional Animal Care and Use Committees of Cleveland Clinic-Florida Research and Innovation Center (CC-FRIC) and the University of Tennessee Health Science Center (UTHSC).

The mouse-adapted SARS-CoV-2-N501Y_MA30_ was provided by Drs. Stanley Perlman and Paul McCray at the University of Iowa, USA (27). The virus was propagated in Calu-3 cells and tittered by plaque assay in VeroE6 cells. Calu-3 cells were maintained in minimal essential medium (MEM) supplemented with 20% fetal bovine serum (FBS), 0.1 mM nonessential amino acids (NEAA), 1 mM sodium pyruvate, 2 mM l-glutamine, 1% penicillin and streptomycin, and 0.15% NaHCO_3_ at 37°C in 5% CO_2_. Vero E6 cells were grown in Dulbecco’s modified Eagle medium (DMEM) supplemented with 10% FBS, 0.1 mM NEAA, 1 mM sodium pyruvate, 2 mM l-glutamine, 1% penicillin and streptomycin, and 0.15% NaHCO_3_ at 37°C in 5% CO_2_.

### Virus infection and titration

The research involving SARS-CoV-2 was conducted within the biosafety level 3 (BSL3) Laboratory at CC-FRIC. Balb/c or C57BL/6 mice were gently anesthetized using isoflurane and subsequently intranasally infected with 10^4^ FPU of SARS-CoV-2-N501Y_MA30_. Post-infection, daily monitoring and weight measurements of the mice were conducted. Tissues were aseptically collected and dissociated in PBS using disposable homogenizers. The viral preparation or supernatants from lung or brain tissue homogenates were subject to sequential dilution in DMEM. These diluted samples were then introduced to VeroE6 cells in 12-well plates to conduct plaque assays (29). After one hour of incubation, the viral inoculums were removed, and the cells were overlaid with a 1.2% agarose solution supplemented with 4% FBS. After 3-day incubation, the cells were fixed with formaldehyde, and the overlays were meticulously eliminated, facilitating visualization of the resulting plaques through the application of a 0.1% crystal violet stain.

### Behavioral experiments

C57BL/6 mice aged 6-9 weeks were utilized to investigate behavioral responses using a battery of tests as described in **Fig. 2**. We conducted a well-established behavioral test battery in mice fourteen days after intranasal administration of SARS2-N501Y_MA30_ (10^4^ PFU/ mouse). See detailed descriptions below.

### Anxiety-like behaviors

#### Open Field Test

Assessed exploratory and anxiety behaviors in an open-field box for 6 minutes. The Open Field task was used to determine the general activity levels, gross locomotor activity, exploration habits of the mice, and anxiety-like behaviors. The evaluation occurred within a cubic, white plastic enclosure measuring 30 cm × 30 cm × 30 cm. Mice were introduced to a corner of the arena and given 6 minutes to acclimate to the unfamiliar surroundings. Their activities were recorded via video. Subsequently, the recorded footage, encompassing metrics such as distance traversed, velocity, resting intervals, movement duration, and time spent in both the center and corners of the arena, was analyzed utilizing Anymaze software. This examination serves as an initial assessment of locomotor activity and anxiety-related behaviors, particularly focusing on the duration spent in the central area.

#### Elevated Plus Maze Test

Evaluated anxiety-like behavior by recording entries into and time spent in the open arms of a maze for 6 minutes. It was conducted as previously described (30). The EPM apparatus, 40 cm high from the floor, consists of two open arms (35 × 10 cm) and two closed arms (35 × 10 cm), those two parts stretch perpendicular to each other and connect in a center platform (5 cm). Mice were placed in the center zone facing an open arm and allowed to explore the maze freely for 6 minutes. The time spent in open arms, closed arms, and behaviors of head dipping, sniffing, and grooming were analyzed.

### Anxiety-like behaviors

#### Tail Suspension Test

Conduct the tail suspension test to assess depression-like behaviors. Suspend the mice by their tails for 6 minutes using adhesive tape in a controlled environment. Record and analyze the duration of immobility during the predetermined time period (6 minutes). The state of immobility is defined by the absence of movement. The detection of immobility time in these tests serves as a behavioral measure to evaluate the impact of various interventions, such as drug treatments or genetic manipulations, on the animal’s response to stress or despair.

#### Forced Swimming Test

Conduct the forced swimming test to assess depression-like behavior. Place the mice individually in a water-filled container for 6 minutes and record their swimming and immobility behaviors. Manually analyze the behavioral results by determining the immobility time for each mouse. Compile the data and calculate the mean immobility time for each experimental group.

### Bulk RNA sequencing

C57BL/6J mice were intranasally (i.n.) infected with 10^4^ FPU of SARS-CoV-2-N501Y_MA30_, as described previously. At specified time points post-infection, whole brains were harvested for RNA extraction using the RNeasy Lipid Tissue Mini Kit (74804, Qiagen, Hilden, Germany). RNA concentration and purity were determined by NanoDrop 2000 spectrophotometer (Thermo Fisher Scientific, Wilmington, DE, USA). Library preparation was performed with the Illumina Stranded mRNA Prep Kit (20040532, Illumina, San Diego, CA, USA), adhering to the manufacturer’s instructions. Sequencing was executed on the DNBSEQ-G400 platform (Innomics Inc. Cambridge, MA, USA). Data analysis and visualization was conducted on the Dr. Tom online analysis platform, utilizing the following pipeline: 1. Data Cleaning: Sequencing data was filtered with SOAPnuke, yielding clean reads stored in FASTQ format. 2. Alignment: Clean reads were aligned to the reference genome using HISAT2. 3. Fusion Gene Detection: Ericscript (0.5.5-5) was employed to identify fusion genes. 4. Differential Splicing Gene (DSG) Analysis: rMATS (v4.1.1) was utilized for detecting DSGs. 5. Gene Set Alignment: Bowtie2 aligned the clean reads to the gene set. 6. Expression Quantification: RSEM (v1.2.28) calculated gene expression levels, providing read count, FPKM, and TPM metrics.7. Differential Expression Gene (DEG) Analysis: DESeq2 conducted DEG analysis, with a significance threshold set at Q value ≤ 0.05. 8.

Visualization: A heatmap of DEGs was generated using Pheatmap, based on the DEG analysis results. 9. Phenotypic Insight: GO (Gene Ontology, http://www.geneontology.org/) and KEGG (Kyoto Encyclopedia of Genes and Genomes, https://www.kegg.jp/) enrichment analyses of annotated DEGs were performed using Phyper, based on the Hypergeometric test. Significance levels of terms and pathways were adjusted by Q value, with a stringent cutoff (Q value ≤ 0.05).

### Quantitative real-time PCR analysis of viral RNA

Total cellular RNA was isolated using the Direct-zol RNA miniprep kit (Zymo Research, Irvine, CA) following the manufacturer’s protocol including a DNase treatment step.

Total RNA (200 ng) was used as the template for first-strand cDNA synthesis. The resulting cDNA was used to quantify the SARS-CoV-2 RNA levels by real-time quantitative PCR using Power SYBR green PCR master mix (Applied Biosystems, Waltham, MA). Average values from duplicates of each sample were used to calculate the viral RNA level relative to the GAPDH gene and presented as 2−ΔCT, as indicated (where CT is the threshold cycle). CT values of gRNA and sgRNA from uninfected mice (0 dpi) are constantly > 35. The sequences of the primers used are listed in the **S1 Table**.

### Brain slice preparation, S1 protein perfusion, and fEPSP recordings

Mice were euthanized with overdosed isoflurane and whole brains were dissected into pre-oxygenated (5% CO_2_ and 95% O_2_) ice-cold high sucrose dissection solution containing (in mM): 205 sucrose, 5 KCl, 1.25 NaH_2_PO_4_, 5 MgSO_4_, 26 NaHCO_3_, 1 CaCl_2_, and 25 Glucose and sliced into 300 µm on a Leica VT1000S vibratome (31). The transverse amygdala slices were then transferred into the normal artificial cerebrospinal fluid (ACSF) containing (in mM): 115 NaCl, 2.5 KCl, 2 CaCl_2_, 1 MgCl_2_, 1.25 NaH_2_PO_4_, 11 glucose, 25 NaHCO_3_, bubbled with 95% O_2_/5% CO_2_, pH 7.35 at 20°C-22°C. Slices were incubated in the ACSF at least 1 hour before recording. Individual slices were transferred to a submersion-recording chamber and were continuously perfused with the 5% CO_2_/95% O_2_ solution (∼3.0 ml/min) at room temperature (20°C - 22°C). The spike protein (BPSbIoscience, #510333) was diluted to 167 ng/ml and perfused to the brain slices in the ACSF. For the field EPSP (fEPSP) experiments, neurons were held in current-clamp mode with a pipette solution containing (in mM): 2 KCl (mOsm = 290, adjusted to pH 7.25 with KOH). A concentric bipolar stimulating electrode (FHC Inc., Bowdoin, ME) was positioned in the cortical inputs in the amygdala slices. Away from the stimulating electrode around 400 μm is a glass recording electrode. EPSPs were recorded in current-clamp mode every 20 seconds and continuously recorded the EPSPs for at least 1 hour. Data were acquired at 10 kHz using Multiclamp 700B and pClamp 10.

### ATP measurement

Brains were removed from the skull and dissected. Brain slices were then placed in 200 µl ice-cold PBS, in the presence of vehicle or spike protein (167 ng/ml, BPS BIoscience, #510333) for the indicated time (10, 30, 60 minutes). To quantify ATP released from the brain slice, we employed an ATP Determination kit (Invitrogen, A22066). Briefly, a 100 µl reaction mixture was added to the 96-well cell culture plate, which contains a 10 µl sample or standard solution. After 15 minutes of incubation in the dark, the plate was read using a Synergy Neo2 hybrid multimode microplate reader (BioTek, Winooski, VT, USA). ATP concentrations were determined by reference to a standard curve.

### Histology

Tissues (lungs, brain) were collected and fixed in zinc formalin. Following fixation, the lungs were processed for paraffin embedding and sliced into a 4 μm section and the brain was sectioned into 30 µm by a vibratome for subsequent hematoxylin and eosin (H&E) staining by Immunohistochemistry Core of Cleveland Clinic Lerner Research Institute and Immunohistochemistry Core at the University of Tennessee Health Science Center. We have used two serial lung sections (six fields/section) from each animal and a total of 4 to 5 animals per group. Acute lung injury severity was evaluated with ATS guidelines (32) for neutrophil infiltration in alveolar and interstitial space, hyaline membranes, alveolar wall thickening, and proteinaceous debris deposition. Briefly, a scoring system (0–2) was employed for each of the criteria mentioned. An average score of 0 indicated absence of injury, 1 indicated mild to moderate injury, and 2 indicated severe injury.

### Immunohistochemistry

Following the behavioral procedures indicated in the text and figures, the mice were euthanized with overdosed isoflurane and were fixed in Zinc Formalin. Following fixation, we used a vibratome (Leica VT-1000S) to dissect 30 µm amygdala coronal slices, which were collected in ice-cold PBS. To complete immunofluorescence staining, slices were placed in Superblock solution (Thermo Fisher Scientific) plus 0.2% Triton X-100 for 1 hour and incubated with primary antibodies (1:1000 dilution) at 4°C for 24 hours (31). Primary antibodies we used include mouse monoclonal antibody to dsRNA (Millipore Cat# MABE1134); Rabbit polyclonal to Iba1 (Abcam Cat# ab108539); Rabbit polyclonal to GFAP (Abcam Cat# ab7260) and mouse anti-NeuN (Cell Signaling Cat# 24307). We then washed and incubated slices for one hour with secondary antibodies diluted at a ratio of 1:200 (Alexa Fluor 488 goat anti-rabbit IgG (H+L) (Thermo Fisher Scientific Cat# A-11008); Alexa Fluor 594 goat anti-mouse IgG (H+L) (Thermo Fisher Scientific Cat# A-11032); Alexa Fluor 647 goat anti-mouse IgG (H+L) (Thermo Fisher Scientific Cat# A-21235). VectaShield H-1500 (Vector Laboratories Cat# H-1500) was used to mount slices, while regular fluorescent DIC microscopy and confocal microscopy were used to image the slices.

For SARS-CoV-2 antigen detection, tissue slides underwent a series of incubation steps. Initially, slides were treated with a blocking reagent (10% normal goat serum) for 30 minutes to reduce non-specific binding. Subsequently, a rabbit polyclonal antibody against the SARS-CoV-2 nucleocapsid protein (dilution: 1:4,000, catalog number: 40143-T62, Sino Biological) was applied to the slides for 15 minutes. Following the primary antibody incubation, slides were further incubated with Rabbit Envision (Dako) and diaminobenzidine (Dako) as a chromogen to visualize the antigen-antibody complexes.

### Dendrites and spine morphologic analyses

After their behavioral testing, we measured the density of dendritic spines on amygdala neurons by using Golgi staining (n = 5 per group). Mice brains were collected, fixed, and processed for the Golgi staining according to the protocol provided by the FD Rapid Golgi Stain Kit (PK401, FD NeuroTechnologies). All images were deconvolved within the Zeiss Application Suite software. The number of dendritic spines was analyzed using plug-in SpineJ as described (33), with modifications. Spines were examined on dendrites of amygdala neurons that satisfied the following criteria: (1) presence of untruncated dendrites, (2) dark and consistent Golgi staining throughout all dendrites, and (3) visual separability from neighboring neurons. We counted the number of dendritic spines along the second dendritic branch at distances from 50 to 100 µm from the soma, in images obtained at 630× magnification. For each neuron, three to five dendritic segments 10 µm in length were analyzed. For each group, 6-10 neurons/mice were analyzed. We used ImageJ software to analyze spines. The analysis of dendritic spines includes the number of spines and spine density, which are critical indicators of synaptic function (34, 35). Spines were classified as thin if they had a long, slender neck and small, Spine length > 0.5 µm; Head perimeter = 2 μm to 3 μm; mushroom, if they had a well-defined, thick neck and spine length > 0.5 µm; Head perimeter ≥ 3 µm; or stubby, if they were short and thick, without a distinguishable neck, spine length ≤ 0.5 µm; or filopodia if they were long and curved, spine length > 0.5 µm, head perimeter ≤ 2 μm.

### Statistical analysis

One-way ANOVA and Tukey’s post-hoc multiple comparison tests were used for statistical comparison of groups. An unpaired Student’s t-test was used to compare results between the two groups. p < 0.05 was considered statistically significant, and we did not exclude potential outliers from our data except the ones that did not receive successful aversive conditioning. The graphing and statistical analysis software GraphPad Prism 8 was used to analyze statistical data, which was presented as means ± SEM. Sample sizes (n) are indicated in the figure legends, and data are reported as biological replicates (data from different mice, and different brain slices). Each group contained tissues pooled from 4-5 mice. Due to variable behavior within groups, we used sample sizes of 10-16 mice per experimental group as we previously described in earlier experiments (31). In behavioral studies, we typically studied groups with five randomly assigned animals per group. The experiments were repeated with another set of four animals until we reached the target number of experimental mice per group.

## RESULTS

### Neuroinvasion following Respiratory Infection of SARS-CoV-2

The inherent low affinity of the ancestral viral spike (S) glycoprotein for mouse ACE2 (mACE2) renders mice naturally resistant to SARS-CoV-2 infection (22, 23). However, specific mutations in the viral spike protein, such as Q498Y, P499T, and N501Y, can enhance binding to mACE2, resulting in asymptomatic infection in mice (36). Through serial passages of a recombinant SARS-CoV-2/spike N501Y virus in BALB/c mice, we successfully isolated a highly virulent mouse-adapted SARS-CoV-2 strain, designated as SARS2-N501Y_MA30_ (27). Interestingly, young C57BL/6 mice exhibited reduced sensitivity to symptomatic infection by SARS2-N501Y_MA30_ (**Fig. 1A**), making them an optimal model for studying the long-term repercussions of COVID-19. Thus, we intranasally infected mice with 10^4^ PFU/mouse of SARS2-N501Y_MA30_ (27). Comparable to our previous findings, young Balb/c mice developed lethal disease, whereas young C57BL/6 mice only displayed minimal weight loss of less than 20% and swift recovery within approximately one week (**Fig. 1B, C**). In C57BL/6 mice, viral titers peaked at 2 days post infection (dpi) in the lungs, followed by the presence of viral titers in the brain peaking at 4 dpi (**Fig. 1D, E**). Consistent with the observation, both viral genomic RNA (gRNA) and subgenomic RNA (sgRNA) were identified in the lung **(Supplemental Fig.1**) and brain (**Fig. 1F, G**). Immunofluorescence targeting double-stranded RNA (dsRNA) further validated virus RNA replication presence in amygdala brain slices at 4 dpi, which subsequently disappeared by 14 dpi. (**Fig. 1H-J**), with similar observations in the prefrontal cortex (PFC), albeit with slight distinctions (**Fig. 1K**). These results collectively suggest direct viral infiltration into the brain, alongside respiratory infection. Remarkably, the production of infectious SARS-CoV-2 appeared to be controlled in the brain, as the titer is extremely low (**Fig 1E**) and it did not induce major pathological changes within brain tissues **(Supplemental Fig. 2)**. Further investigation revealed that SARS-CoV-2 replication was predominant in neuron cells in the amygdala, rather than in microglia or astrocytes (**Fig. 1L-N**). These findings indicated a SARS2-N501Y_MA30_ neuroinvasion with a peak at 4 dpi followed by a quick clearance.

**Figure. 1.**
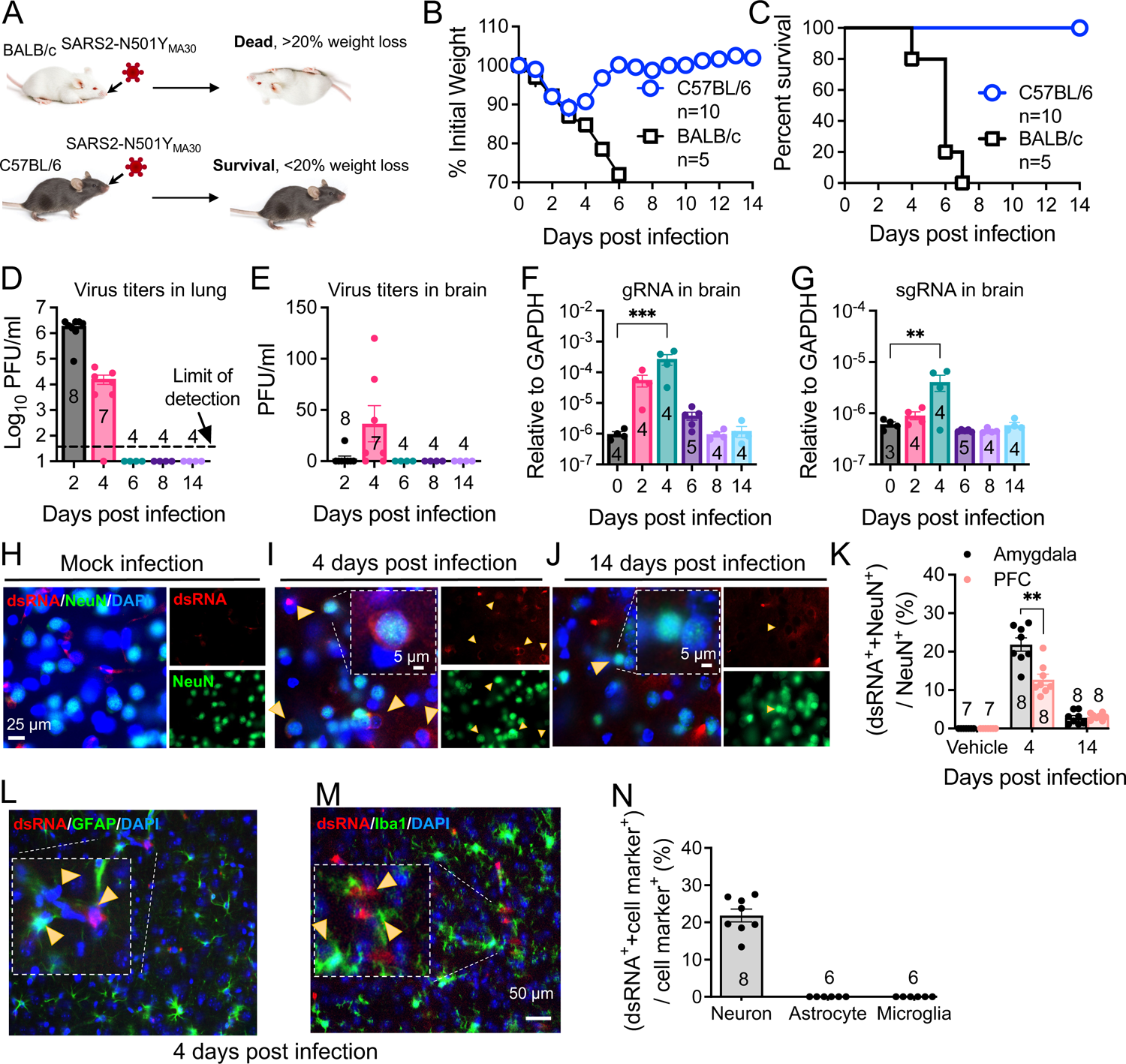
Outcomes of Intranasal Infection with SARS2-N501Y_MA30_. **(A)** Schematic depicting the outcomes of infection in young BALB/c and C57BL/6 mice following administration of 10^4^ PFU of SARS2-N501Y_MA30_. **(B-C)** Daily monitoring of body weight (B) and survival (C) in young BALB/c and C57BL/6 mice post-infection. **(D-E)** Virus titers in the lungs (D) and brains (E) of C57BL/6 mice infected with SARS2-N501Y_MA30_ at the indicated days post-infection. **(F-G)** Viral genomic RNA (gRNA) (F) and subgenomic RNA (sgRNA) (G) levels in brain tissues from SARS2-N501Y_MA30_ infected C57BL/6 mice. The levels of viral gRNA and sgRNA were normalized to GAPDH and presented as 2^-ΔCT (n = 4 or 5 mice per group). CT values for viral genomic RNA (gRNA) or subgenomic RNA (sgRNA) from mock-infected tissues were consistently greater than 35. Statistical significance: ***p = 0.001, **p = 0.0017, determined by ordinary one-way ANOVA. **(H-J)** Immunofluorescence staining targeting dsRNA (red), neuronal nuclear protein (NeuN, green), and nuclei (DAPI, blue) in amygdala brain slices collected from mice infected with SARS2-N501Y_MA30_ infection at 4 or 14 days post (I-J), compared to mock infection (H). Arrows indicate dsRNA and NeuN-positive neurons. Inserts show enlarged single-cell images. **(K)** Percentage of dsRNA-positive cells (dsRNA^+^) in slices from the amygdala and the prefrontal cortex (PFC). The peak of dsRNA^+^ cells is observed at 4 dpi. Statistical significance: **p = 0.0012, determined by a two-tailed unpaired Student’s t-test. **(L-M)** Immunofluorescence targeting dsRNA (red), astrocyte (glial fibrillary acidic protein, GFAP, green) or microglia (Ionized calcium-binding adaptor molecule 1, Iba1, green), and nuclei (DAPI, blue) in amygdala slices from infected mice. **(N)** Comparison of dsRNA^+^ cells with NeuN^+^, GFAP^+^, and Iba1^+^ at 4 dpi.. Data are presented as mean ± SEM.

Histological analysis of lung samples revealed acute lung injury using American Thoracic Society (ATS) guidelines for experimental acute lung injury (ALI) based on neutrophil infiltration in alveolar and interstitial space, hyaline membranes, alveolar wall thickening, and proteinaceous debris deposition (32). SARS-CoV-2 infection induced mild lung injury which peaked on day 6, accompanied by viral antigen presence at 2 dpi and disappeared after 6 dpi **(Supplemental Fig 3)**. However, these changes, including infiltration and viral presence, resolved after a 14-week period. To explore host defense responses in SARS-CoV-2-infected mice, we assessed cytokine/chemokine gene expression in the lungs **(Supplemental Fig 1)**. Notably, these genes exhibited significant upregulation at 2 dpi or 4 dpi, gradually subsiding to baseline levels after 6 dpi. These findings collectively illustrate a mild respiratory disease in the infected mice, which subsequently resolved within 14 days post-infection.

### SARS2-N501Y_MA30_ infection induces significant changes in anxiety- and depression-like behaviors

Our investigation successfully detected viral dsRNA within the amygdala (**Fig. 1K**), an area pivotal in coordinating defensive responses in mice (37) and emotional behaviors in humans (28). This finding prompted us to postulate that the infection may still exert influence over the functionality of this region and consequently modify the defensive behaviors of infected mice. To evaluate the impact of SARS-CoV-2 infection on anxiety- and depression-like behaviors, we conducted a well-established behavioral test battery in mice fourteen days after intranasal administration of SARS2-N501Y_MA30_ (10^4^ PFU/ mouse) (**Fig. 2A**). We chose the 14-day time point for behavioral testing since viral infections were observed to have cleared by then. C57BL/6 mice typically recovered from symptomatic infection at 10^4^ PFU/mouse within about a week (**Fig. 1B**). Additional indicators, such as body temperature, fur condition, daily food, and water intake over time (daily records), and mouse locomotion behavior, were similar to those of mock-infected mice at 14 dpi.

**Figure. 2.**
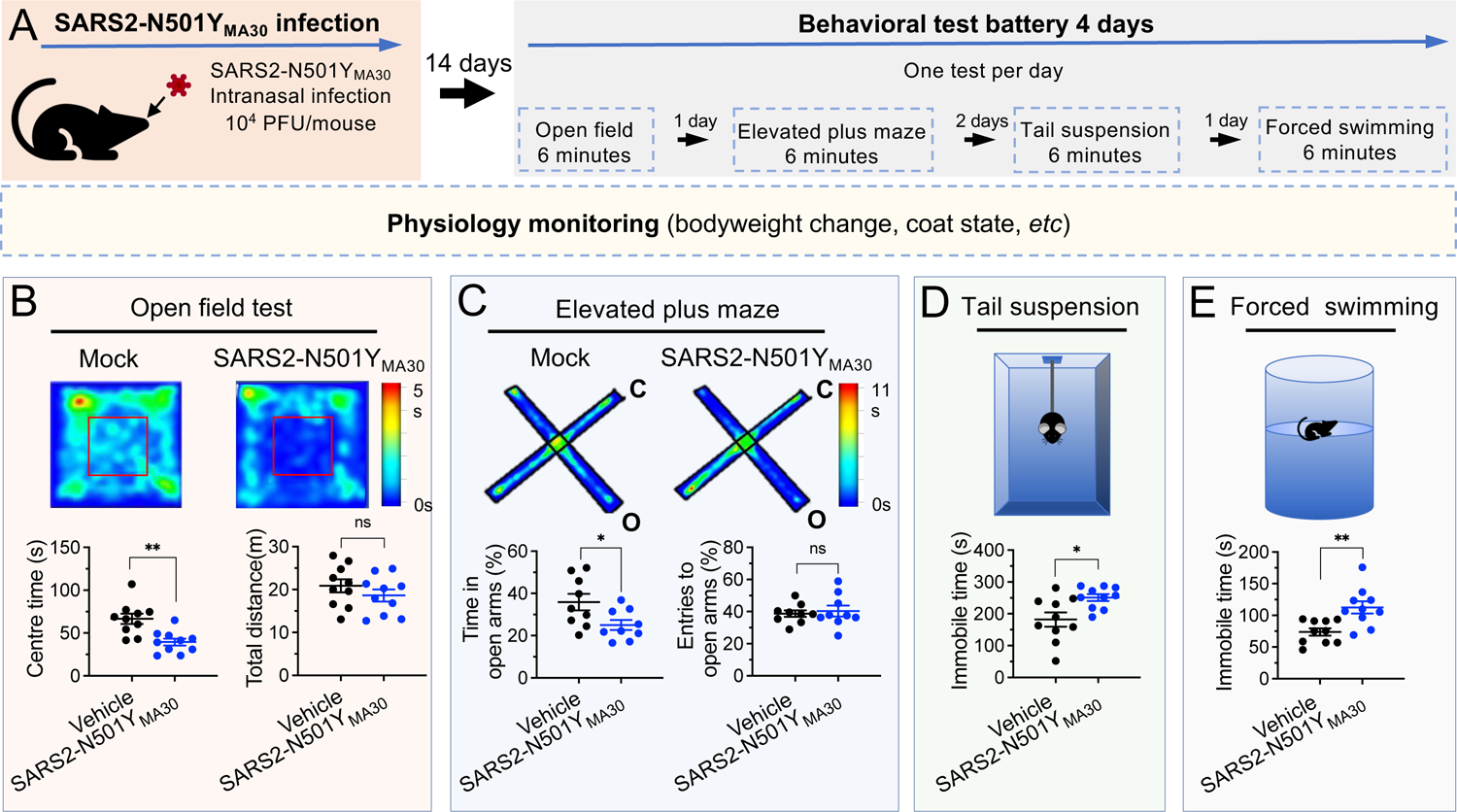
Induction of anxiety- and depression-like Behaviors in Mice Following SARS2-N501Y_MA30_ infection. **(A)** Experimental design and timeline illustrating the administration of SARS2-N501Y_MA30_ and the behavioral test battery. **(B)** Open field test. Upper: Representative heat map tracking of activity in the mock and SARS2-N501Y_MA30_ infected mice. Lower: Results depicting the total travel distance and time spent in the center area. Statistical significance: **p = 0.0014 (total travel distance); ns, non-significant, p = 0.2927 (center area). **(C)** Elevated plus maze test. Upper: Representative heat map tracking of activity in mock and SARS2-N501Y_MA30_ infected mice in closed arms (C) and open arms (O). Lower: Results indicating the time spent in open arms and the number of entries into open arms. Statistical significance: *p = 0.0299 (time spent in open arms); ns, non-significant, p = 0.6897 (number of entries). **(D-E)** Schematics of the tail suspension (D) and forced swimming (E) tests and results showing immobile time. Statistical significance: *p = 0.0118 (tail suspension test); **p = 0.0029 (forced swimming test). All statistical analyses were performed using a two-tailed unpaired Student’s t-test. The analysis includes data from 10 mice in each group. Data are presented as mean ± SEM.

In the open-field test, the infected mice exhibited a decreased duration spent in the center area compared to the mock infection group, indicative of heightened anxiety-like behaviors (**Fig. 2B, left)**. Remarkably, both the SARS2-N501Y_MA30_ and mock infection groups demonstrated similar locomotor activity, implying a complete recovery of locomotion post-infection (**Fig. 2B, right)**. In the elevated plus-maze test, mice in the infected group significantly reduced their time spent in the open arms (**Fig. 2C, left)**, while the number of entries into the open arms remained unaltered (**Fig. 2C, right)**, further substantiating an augmentation in anxiety-like behaviors. Consistent outcomes were observed in the tail suspension and forced swimming tests, both are widely used for evaluating depression-like behaviors, where mice in the virus-infected groups displayed increased immobile time compared to the vehicle groups (**Fig. 2D, E**). These findings substantiate our conclusion that SARS2-N501Y_MA30_ infection leads to an escalation in anxiety- and depression-like behaviors in mice, indicating potential repercussions on brain function that warrant further exploration.

### Temporal Gene Signatures in the Brain after SARS-CoV-2 Infection

To further elucidate the molecular mechanisms associated with the observed neuroinvasion and subsequent behavioral alterations, we investigated the temporal dynamics of host transcriptional responses in the brain. This was achieved through bulk RNA sequencing (RNA-seq) on total brain RNA from both mock-infected animal cohorts (0 dpi) and animals infected with SARS2-N501Y_MA30_ at 4 dpi, and 14 dpi, respectively (**Fig. 3 and Supplemental Fig. 4)**. Principal component analysis (PCA) unveiled distinct differences among mock and infected groups with samples from both 4 and 14 dpi **(Supplemental Fig. 4A).** The volcano map of differentially expressed genes (DEGs) revealed significant changes in host genes compared to mock-infected animals or between 4 and 14 dpi based on |log2fold| ≥ 1.5, Q value ≤ 0.05 **(Supplemental Fig. 4B)**. In the 4 dpi vs 0 dpi group, 124 genes were upregulated and 6 were downregulated, while in the 14 dpi vs 0 dpi group, 29 genes were upregulated and 2 were downregulated (**Fig. 3A**). There were only 8 genes upregulated and 5 downregulated between 4 and 14 dpi. Notably, 29 upregulated genes in the 14 dpi group also appeared in the 4 dpi group, many of which were associated with synaptic transmission and neuronal development (**Fig. 3B and Supplemental Fig. 4).** Overall, there were 140 genes significantly changed **(Supplemental Fig. 4).**

**Figure. 3.**
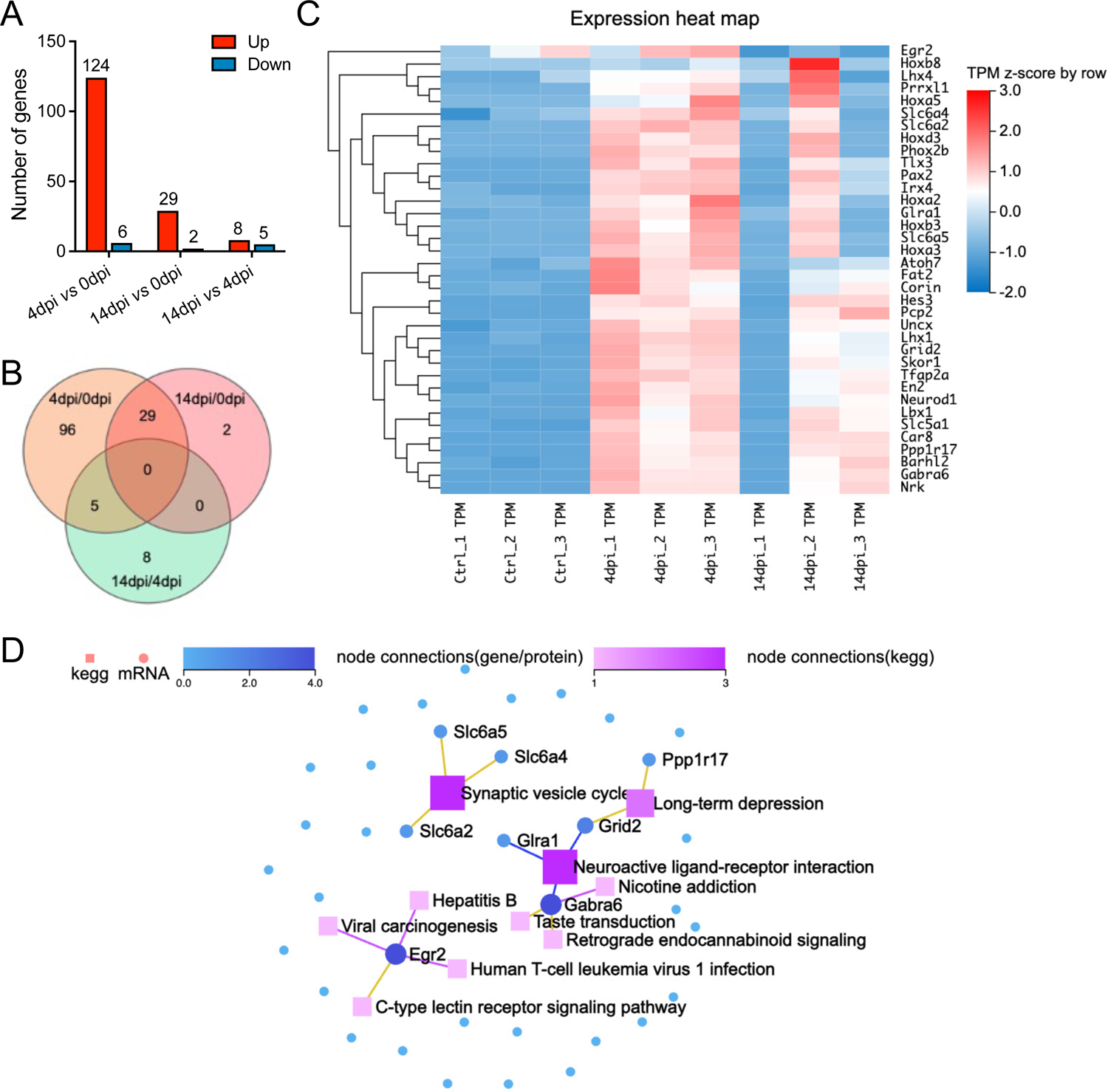
RNA sequencing reveals distinct alterations in the brains of SARS2-N501Y_MA30_ Infected Mice. **(A)** Histogram represents the corresponding numbers of differentially expressed genes (DEGs) between groups. (|log2fold| ≥ 1.5, Q value ≤ 0.05) **(B)** Venn diagram illustrating the differential gene expression analysis between groups of gene sets. The intersections show shared gene expression patterns, while non-overlapping regions indicate uniquely expressed genes. The numbers within each section denote the count of genes. **(C)** Heatmap displaying hierarchical clustering of 36 selected genes associated with neuron functions. **(D)** KEGG Network analysis of key driver genes and enriched KEGG pathways among the 36 selected genes. Dots symbolize genes, squares represent KEGG pathways, and lines connecting genes to squares signify gene enrichment within the respective pathway.

We performed pathway analysis based on the Kyoto Encyclopedia of Genes and Genomes (KEGG; http://www.genome.jp/kegg/) and Gene Ontology (GO) based enrichment analysis on the 140 genes **(Supplementary** Fig. 4C, E). Among them, 36 genes were implicated to be involved in different neuronal functions in the brain (**Fig. 3C),** such as synaptic transmission, including Car8, Gabra6, Pcp2, Slc5a1, Tfap2a, or neuronal development, such as Barhl2, Hoxa5, Ppp1r17, Skor1, Fat2, and Nrk. We conducted a KEGG network analysis to delineate the interconnections among the 36 identified genes and identified genes related to synaptic vesicle cycle (slc6a2, slc6a4, and slc6a5), long-term depression (Ppp1r17 and Grid2), and neuroactive ligand-receptor interaction (Grid2, Glra1, and Gabra6) (**Fig. 3D**). In conclusion, our comprehensive RNA-seq analysis has unveiled significant changes in host gene expression following viral infection. These alterations suggest a reprogramming of neuronal transcriptional states, corroborating the observed behavioral modifications. Moreover, the data indicate a profound influence on synaptic function and neurodevelopmental processes, underscoring the far-reaching effects of SARS-CoV-2 on the central nervous system.

### SARS2-N501Y_MA30_ infection induces spine remodeling in the amygdala

To complement the gene expression changes revealed by our RNA-seq analysis, we next examined the morphological alterations in neurons within the amygdala, a region implicated in depression-related behaviors. This investigation aimed to confirm the impact of SARS2-N501Y_MA30_ infection on synaptic plasticity through the observation of dendritic spine remodeling. Dendritic morphology has been widely implicated in the mechanisms governing synaptic plasticity (35, 38, 39). Dendritic spines constitute the principal target of neurotransmission input within the CNS (40), and their density and structure form the foundation for physiological alterations in synaptic efficacy that underlie learning and memory processes (41). We hypothesized that dendritic structure and plasticity undergo alterations following SARS2-N501Y_MA30_ infection. Fourteen days after SARS2-N501Y_MA30_ infection, brain slices containing the amygdala were meticulously dissected, fixed, and subjected to Golgi staining to label the spines (**Fig. 4A**). Neurons were randomly selected from within the basolateral amygdala, with a focus on secondary branches located 50-100 µm away from the soma (**Fig. 4B, C**). Subsequently, spine morphology was manually visualized and analyzed (**Fig. 4D**).

**Figure. 4.**
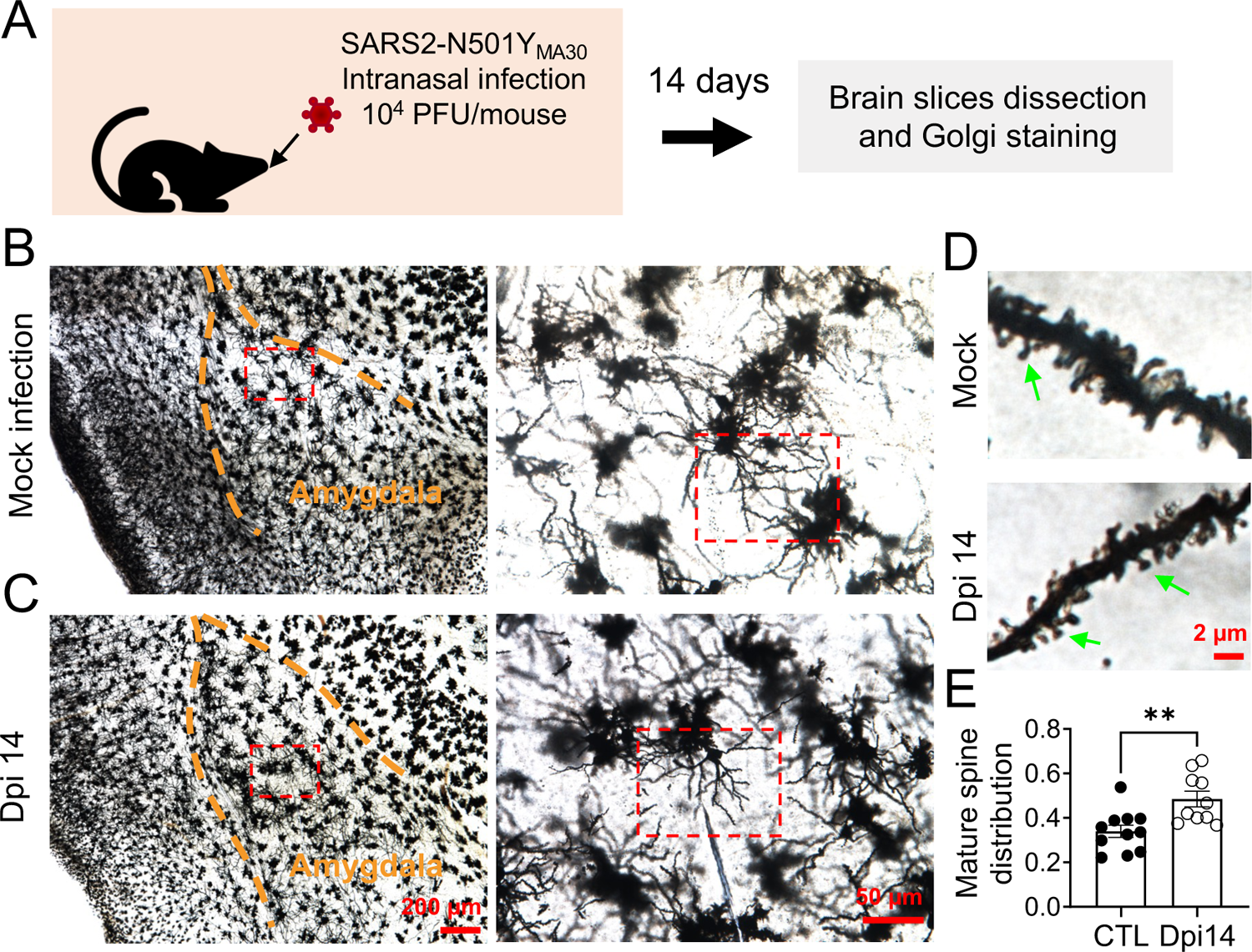
Golgi-Cox Staining of Neurons in the Amygdala of SARS2-N501Y_MA30_ Infected Mice. **(A)** Experimental design and timeline depicting the administration of SARS2-N501Y_MA30_, dissection of brain slices, and Golgi staining process. **(B-C)** Left: Representative Golgi-Cox staining images of amygdala slices from mock-infected (B) and mice infected with SARS2-N501Y_MA30_ at 14 days post-infection (dpi) (C). Scale bar = 200 μm. Right: Enlarged images from the red boxes in the left images for each respective group. Scale bar = 50 μm. **(D)** Representative dendrite images from the red boxes in (B) and (C). Scale bar = 2 μm. Green arrows point out the mushroom spines. **(E)** Comparison of the percentage of mature spines in the mock and SARS2-N501Y_MA30_ infection groups. Statistical significance: **p = 0.0038. The analysis includes data from 11 neurons sourced from 4 mice in each group. Statistical analysis was performed using a two-tailed unpaired Student’s t-test. Data are presented as mean ± SEM.

To characterize spine morphology, we categorized dendritic processes into two distinct morphological classes: mature and immature spines (42). Mature spines, predominantly exhibiting a “mushroom-like” morphology, exhibit more stable postsynaptic structures enriched in α-amino-3-hydroxy-5-methyl-4-isoxazolepropionic acid receptors (AMPARs) and are considered functional spines. Conversely, immature spines, characterized by their thin, stubby, and filopodial features, represent unstable postsynaptic structures with transitional properties. Immature dendritic spines are believed to hold the potential for future synaptic plasticity, either maturing into functional spines or disappearing from the dendrite (43). Spine categories were identified based on parameters described in previous studies and Methods (35, 44).In the brains of mice infected with SARS2-N501Y_MA30_ (14 dpi), we found variations in spine subtypes, notably an elevated proportion of mature spines compared to the mock infection group (**Fig. 4E**). Combined with the gene expression profiles from **Fig. 3** and observations on dendritic morphology, our findings indicate enhanced synaptic plasticity in amygdala neurons fourteen days after SARS2-N501Y_MA30_ infection.

### Spike protein increases microglia-dependent fEPSPs in amygdala slices

Exposure to the SARS-CoV-2 spike protein has emerged as a topic of interest in neurobiology, particularly regarding its impact on neuronal function. Previous research has provided initial insights into how spike protein can influence neuronal morphology and function (45–47). To further explore the mechanistic aspects of this phenomenon, we focused on the amygdala, a region critically involved in emotional processing and defensive behaviors. We perfused amygdala slices from mice with the recombinant B1.351 variant spike protein (167 ng/ml), which has a high affinity to mouse ACE, and monitored changes in fEPSPs for 2 hours (**Fig. 5A**). We observed a significant increase in fEPSPs following exposure to the B1.351 variant spike protein, indicating heightened synaptic transmission and neuronal excitability (**Fig. 5B, C**). Despite the fact that whether SARS-CoV-2 spike protein increases or decreases neuronal activity is controversial (45–48), this observation is noteworthy, as it suggests that the spike protein, when directly interacting with neurons, has the potential to amplify their responsiveness to incoming signals. This heightened neuronal response could, in turn, render individuals more sensitive to various stimuli, potentially leading to alterations in behaviors or physiological reactions. Interestingly, the administration of 50 μM Resveratrol, a known inhibitor of microglial activation (49), effectively counteracted the spike protein-induced enhancement of fEPSPs (**Fig. 5C, D**). This finding suggests that microglia, the intrinsic resident immune sentinels of the brain, may play a pivotal role in mediating the effects of the spike protein on neuronal activity.

**Figure. 5.**
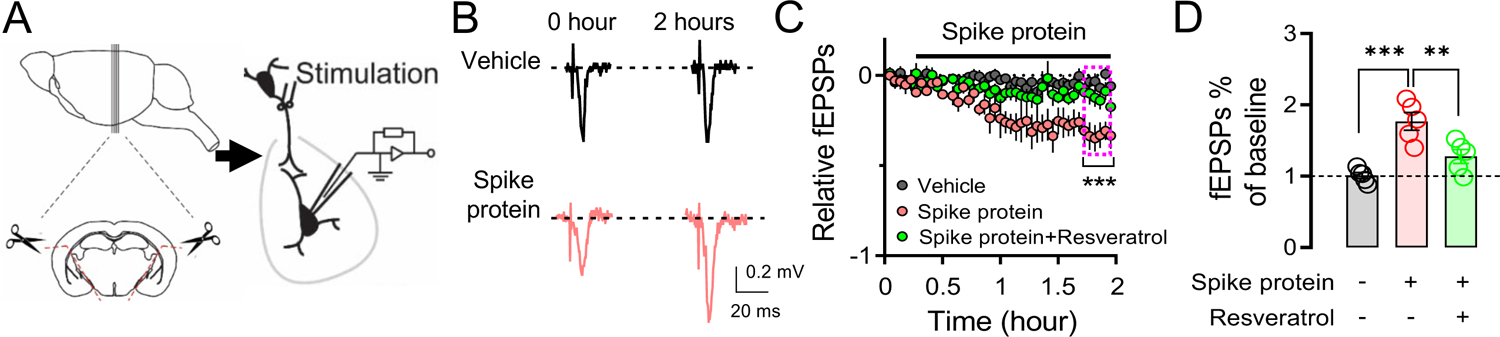
Enhancement of Microglia-Dependent fEPSPs in the Amygdala by SARS-CoV-2 Variant B.1.351 Spike Protein. **(A)** Schematic representation of field excitatory postsynaptic potential (fEPSP) recordings in the Amygadala brain slices. **(B)** Representative fEPSP traces were recorded at the beginning (0 hours) and the end of the 2-hour recordings, during perfusion with either vehicle or spike protein (S1+S2, B.1.351, β variant, BPS Bioscience #510333, 167 ng/ml). **(C)** Average fEPSP data. The spike protein was applied to the slice (167 ng/ml) at the indicated time point (red trace). In the vehicle group, a mock aqueous buffer solution without the spike protein was used (gray trace). The enhancement of fEPSPs by the spike protein was attenuated by pre-treating brain slices with 50 μM Resveratrol (green trace). **(D)** Summarized amplitudes of the last five fEPSPs as shown in (C). Statistical significance: ***p = 0.0003, **p = 0.0082, determined by one-way ANOVA with Tukey’s post hoc multiple comparisons. The analysis includes data from 5 slices in each group. Data are presented as mean ± SEM.

### Microglia activation after SARS-CoV-2 infection or exposure to spike protein

Following the discovery in **Fig. 5**, we conducted further investigations into microglial reactivity during SARS2-N501Y_MA30_ infection. Through immunofluorescence staining for Iba1, we documented a significant activation of microglia, evidenced by their morphological shift from ramified to amoeboid shapes (50). This shift was notable at 4 days post-infection, by 14 days post-infection, microglia activity had not fully reverted to the baseline quiescent state observed at 0 dpi, although it was substantially reduced compared to 4 dpi, as illustrated in **Fig. 6A**. This finding provides further support for the hypothesis that microglial activation may contribute to the observed alterations in neuronal activities within the amygdala following SARS-CoV-2 infection.

**Figure. 6.**
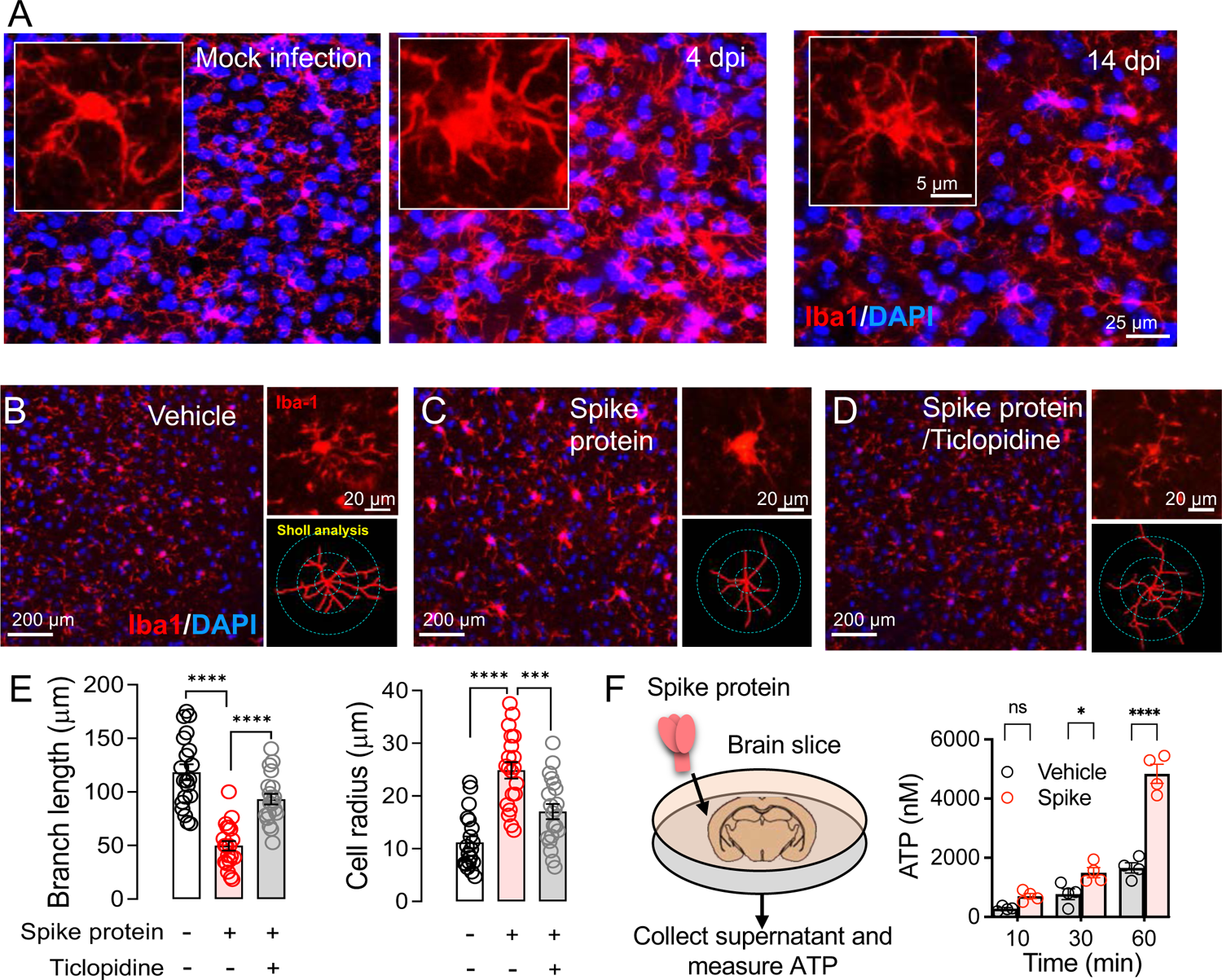
Microglial Activation in Response to SARS-CoV-2 infection and B1.351 Spike Protein. **(A)** Dynamics of Microglial Activation Post-Infection in the Amygdala of Mice. Immunofluorescence staining for Iba1 highlights microglia in amygdala slices from mice at various time points following infection. An enlarged view of individual microglia provided to detail the changes in shape and morphology indicative of activation states. **(B-D)** Microglial response to perfusion with the spike protein. Immunostaining with Iba-1 illustrates microglial morphology in brain slices after a one-hour perfusion with the (B) vehicle or (C) spike protein (S1+S2, B.1.351, β variant, 167 ng/ml), and (D) spike protein following a 30-minute pretreatment with 50 μM Ticlopidine, a P2Y12R antagonist. The series showcases the microglial response to the spike protein, with additional magnified images highlighting the individual microglia and their morphological analysis using Sholl analysis, presented to the right of each panel. **(E)** Comparison of microglial activation assessed by Sholl analysis, which measures the total branch length and radius of cell area. Statistical significance: ****p < 0.0001, ***p = 0.0006, determined by one-way ANOVA with Tukey’s post hoc multiple comparisons. The analysis includes data from 30 cells across 4 slices in each group. **(F)** Quantification of ATP release. Brain slices were incubated in cold PBS with either vehicle or spike protein (167 ng/ml). Supernatants were collected at 10, 30, and 60 minutes post-incubation. The amount of ATP released into the supernatant was quantified. Each dot is an independent brain slice and represents the mean of 2 technical replicates. Statistical significance: *p =0.0374, ****p < 0.0001, determined by two-way ANOVA. Data are presented as mean ± SEM.

Microglia, the brain’s resident immune cells, are highly dynamic, engaging with neurons and other brain cells upon activation. Their processes reach out to neighboring cells, forming a complex network of interactions crucial for understanding the neuroimmune impact of SARS-CoV-2 infection (51). To further elucidate the mechanism by which the spike protein promoted microglia activation, we visualized microglia by immunofluorescence staining for Iba1 in mouse amygdala slices. Our observations confirmed the activation of microglia following SARS-CoV-2 B1.351 spike protein perfusion (**Fig. 6B, C**). Interestingly, the activation was attenuated by the purinoceptor P2Y_12_ (P2Y12R) antagonist ticlopidine (**Fig. 6C, D**), indicating that the P2Y12R signaling pathway mediates this response. P2Y_12_, selectively expressed in microglia (52), serves as the main adenosine triphosphate (ATP) receptor, initialing microglial activation and mobilization in response to ATP, a potent neuronal chemoattractant released from neurons (19, 53). Additionally, it is worth noting that SARS-CoV-2 infection has been shown to induce ATP release from host cells, including neural cells, via specific channels (54, 55), a phenomenon we corroborated by detecting elevated ATP levels in brain slices perfused with spike protein (**Fig. 6E, F**).

### Brain Immune Inflammation and Anti-Viral Responses during Viral Infection

Cytokines and chemokines serve as crucial indicators of anti-viral response and immune inflammation. To delineate the immune responses within the brain during SARS-CoV-2 infection, we conducted quantitative reverse-transcription PCR (RT-qPCR) to profile cytokines and chemokines expression across various time points in the brain (**Fig. 7**). Contrary to expectations, the expression levels of interferon genes, including IFN-β, γ, and λ, as well as the tumor necrosis factor gene, remained unaltered in the infected brain tissues. However, it is noteworthy that transcripts encoding pro-inflammatory cytokines, such as interleukin-1α (IL-1α), interleukin-6 (IL-6), interleukin-8 (IL8/CXCL1), and especially monocyte chemoattractant protein-1 (MCP1/CCL2), exhibited significant upregulation at 2- or 4-days post-infection. These temporal increases coincided with the presence of viral RNA in the brain. Importantly, it is worth highlighting that the alterations observed in immune gene expression within the brain were modest in comparison to those observed in the lungs and returned to baseline levels after 6 days post-infection. This discovery underscores the presence of restrained immune inflammation and anti-viral responses within the brain. The observed changes in immune gene expression warrant further comprehensive investigations to unravel the underlying mechanisms and identify potential targets for therapeutic intervention.

**Figure. 7.**
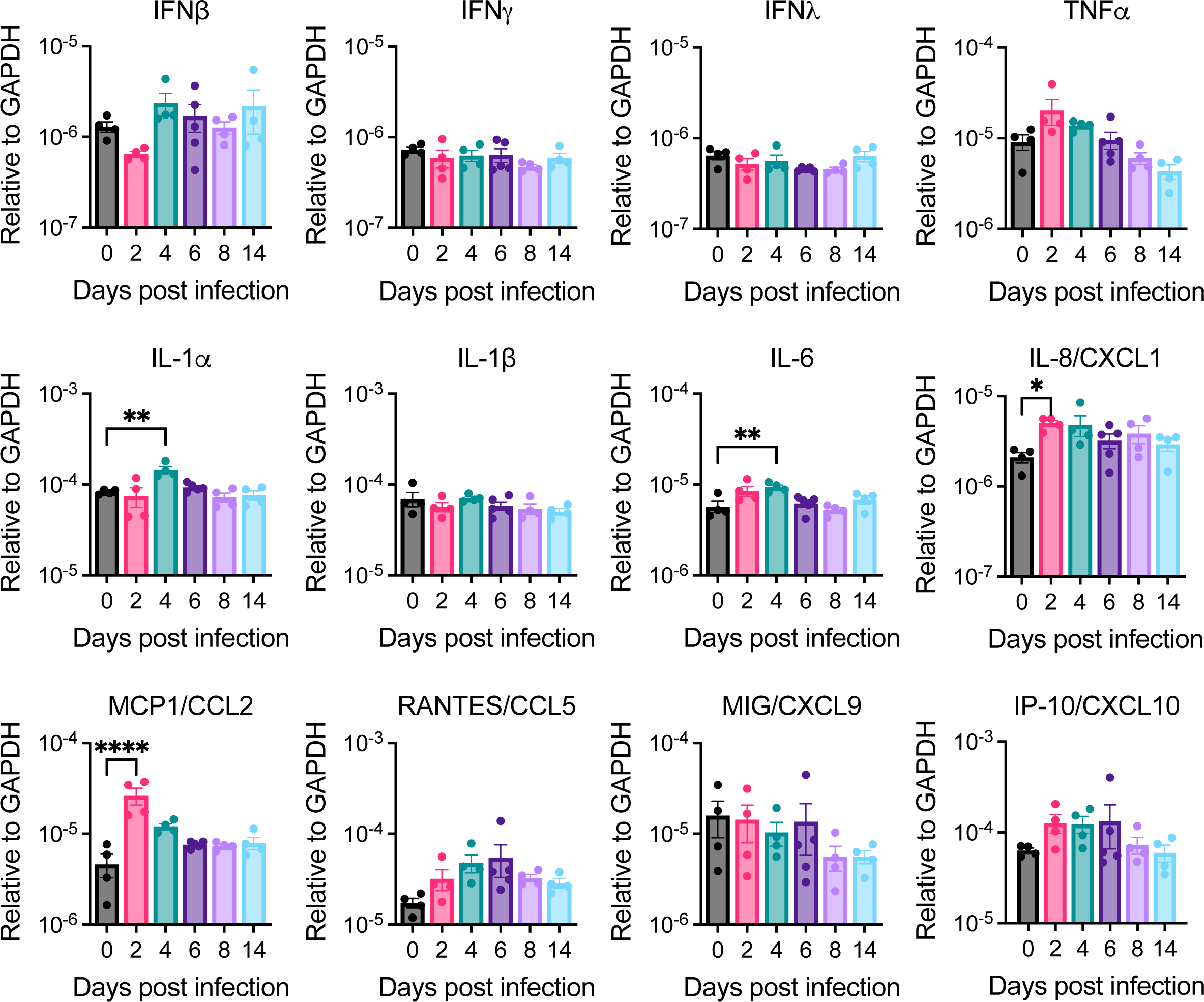
Cytokines and Chemokines Induced in the Brain of SARS-CoV-2-Infected Mice. Brains of C57BL/6 mice intranasally infected with 10^4^ PFU of SARS2-N501Y_MA30_ were harvested at indicated days post-infection. Cytokine and chemokine transcripts were measured by quantitative real-time PCR (qRT-PCR) analyzing total RNA extracted from mock-infected (0 dpi) and infected young C57BL/6 mice. Each brain was collected from one individual mouse. Mock (0 dpi), 2, 4, 8, 14 dpi: n=4; 6 dpi: n=5. The levels of transcripts were normalized to GAPDH and presented as 2^-ΔCT. Statistical significance: *p < 0.05, **p < 0.01, ****p < 0.0001, determined by ordinary one-way ANOVA. Data are presented as mean ± SEM.

## DISCUSSION

### Neurotropism of SARS-CoV-2 and Behavioral Alterations

Our findings revealed the presence of SARS-CoV-2 in the CNS following intranasal inoculation in a SARS2-N501Y_MA30_ infection mouse model (**Fig.1**). These results are consistent with previous reports indicating that SARS-CoV-2 can invade the human brain (5, 6), supporting the notion of neurotropism. SARS-CoV-2 probably enters the central nervous system (CNS) through two potential pathways (56). The first involves accessing the CNS via the neural-mucosal interface in the olfactory mucosa, enabling the virus to spread from the periphery olfactory neurons into the neurons of the olfactory bulb. The second pathway is through entry into the brain via blood circulation, potentially breaching the blood-brain barrier (BBB). In this scenario, the integrity of the BBB could be disrupted by inflammatory responses triggered by the infection.

Notably, the observed viral invasion of the CNS was accompanied by significant alterations in anxiety- and depression-like behaviors, as evidenced by behavioral assays (**Fig. 2**). Anxiety and depression are prevalent neuropsychiatric disorders associated with significant morbidity and mortality worldwide. The link between other viral infections and mental health disturbances has been reported in previous studies, with evidence of viral neuroinvasion contributing to behavioral changes (57, 58). Our study provides valuable evidence supporting a direct association between SARS-CoV-2 neurotropism and anxiety- and depression-like behaviors. These findings underscore the importance of considering the potential neuropsychiatric consequences of COVID-19 and highlight the need for comprehensive mental health assessments and interventions during and after the acute phase of infection.

### Microglial Activation, Neuroinflammation, and Neuronal Activity

Microglia, as the primary immune cells of the CNS, play a critical role in maintaining brain homeostasis and protecting against infections (59). Upon encountering pathogens or damage signals, microglia undergo rapid activation, transforming from a surveillant to an immune-responsive phenotype. In our study, we observed robust microglial activation in brain regions where viral particles were detected, suggesting a potential role for microglia in mediating SARS-CoV-2-induced neurobehavioral alterations. While microglia itself may not be a direct target of SARS-CoV-2 infection (4, 60), the initial invasion of SARS-CoV-2 in the neuron cells of the brain likely triggers their activation.

Microglial activation can lead to neuroinflammation, characterized by the release of pro-inflammatory cytokines, chemokines, and reactive oxygen species (ROS). Chronic neuroinflammation has been implicated in the pathogenesis of various neuropsychiatric disorders, including anxiety and depression (61). Notably, our transcriptomic analysis of SARS-CoV-2-infected brains revealed upregulation of inflammatory and cytokine-related pathways, supporting the hypothesis of microglia-driven neuroinflammation as a mechanism underlying behavioral changes. While our whole-brain qPCR and bulk RNA seq revealed minimal changes in immune responses, it is conceivable that the immune responses induced by SARS-CoV-2 are relatively subdued and localized to specific brain regions. Therefore, the comprehensive assessment of whole-brain studies may underestimate the extent of regional immune responses.

Purinergic receptor P2Y_12_, a G protein-coupled receptor exclusively found in microglia (52), played a pivotal role in the brain’s responses to extracellular nucleotides like ATP and ADP. Upon activation by ATP, microglia not only accumulate but also modulate neuronal activity (16, 17), implicating ATP, known for its swift release from activated neurons (62), as a swift mediator in the neuroimmune response to SARS2-N501Y_MA30_ infection. This activation leads to the release of pro-inflammatory cytokines, which are known to influence neuronal excitability and synaptic transmission (63), potentially exacerbating anxiety- and depression-like behaviors (64). Indeed SARS-CoV-2 infection has been shown to induce ATP release from brain cells, including neural cells, through ATP-release channels (54, 55), reinforcing the link between infection and rapid ATP-mediated microglia response. Clinical and laboratory studies support the idea that elevated cytokines like IL-1β, IL-6, CCL2 and TNF-α associated with the development of anxiety and depression-like behaviors in mice (65–68), underscoring the contribution of neuroinflammation to the behavioral manifestations observed following SARS-CoV-2 infection. Our hypotheses detailing the interactions are visually summarized in **Fig. 8**, providing a clear representation of the underlying mechanisms.

**Figure. 8.**
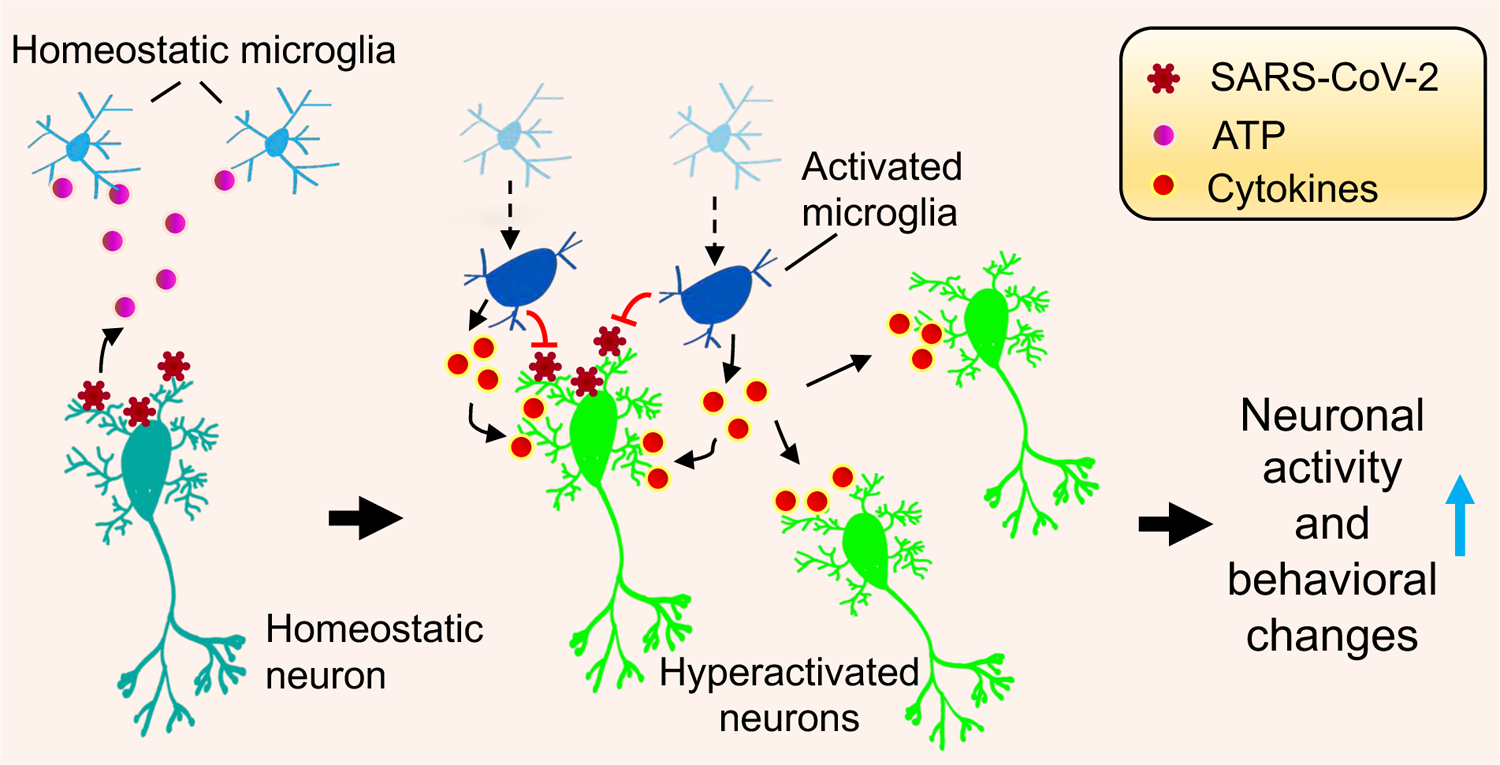
Modeling Neurotropism-induced Behavioral Changes by SARS2-N501Y_MA30_. This schematic illustrates the potential mechanism of SARS-CoV-2 targeting neurons, prompting ATP release, which in turn recruits and activates microglia. The ensuing cascade of microglial response and pro-inflammatory cytokine production is hypothesized to amplify neuronal activity, potentially altering defensive behaviors associated with anxiety and depression.

### Microglial Inhibition and Neuronal Activity Reversal

To elucidate the specific role of microglia in mediating neuronal activity following SARS-CoV-2 infection, we selectively inhibit microglia using a pharmacological approach.

Intriguingly, this inhibition markedly reduced the heightened neuronal activity induced by viral spike protein. This observation indicates a direct involvement of microglia in the development of abnormal neuronal activity in response to SARS-CoV-2 infection. The observed neuronal activity rescue upon microglial inhibition aligns with previous studies showing that microglial activation is a key driver of neuronal activity changes in various CNS disorders (69). Understanding the interactions between microglia and neurons and how SARS-CoV-2 infection influences these processes in the amygdala is crucial to comprehending the neurological consequences of the virus. Such insights could offer valuable information to guide potential interventions or treatments for neurological manifestations observed in long COVID. Further research is needed to fully elucidate the intricate mechanisms underlying microglial activation and its impact on neuronal functions in response to SARS-CoV-2 infection in the brain.

### Inflammatory Pathways and Therapeutic Interventions

Our hypothesis suggests that microglia-driven neuroinflammation contributes to the behavioral changes seen in long COVID-19. This highlights the therapeutic potential of targeting neuroinflammatory pathways to ameliorate anxiety- and depression-like symptoms associated with the disease. The transcriptomic analysis of brains affected by the virus has identified upregulated inflammatory pathways, indicating potential targets for therapeutic interventions. Anti-inflammatory agents and immunomodulatory drugs that selectively dampen microglial activation may offer a promising approach for treating the neuropsychiatric consequences of SARS-CoV-2 infection. However, further research is needed to identify specific inflammatory mediators and downstream signaling pathways responsible for the observed behavioral changes.

### Limitations and Future Directions

While our study provides valuable insights into the link between SARS-CoV-2 neurotropism, microglial activation, neuronal activity, and anxiety- and depression-like behaviors, several limitations should be acknowledged.

First, in our pursuit to unravel the etiology of neurological disorders post-COVID-19, we identified behavioral phenotypes emerging within weeks post-infection. The onset of neurological symptoms post-COIVD-19 can vary greatly among individuals, manifesting during the acute phase, post-recovery, or even several months later. The persistence of PASC symptoms in humans can range from weeks to several months, and in some cases, may extend even further. We acknowledge the differences in lifespan and sensitivity to SARS-CoV-2 between our murine model and human subjects. We chose the 14-day time point for behavioral testing as viral infections were observed to have cleared, and the conditions of infected mice had fully recovered. This rapid recovery in the mouse model confers a significant advantage for our mechanistic studies, eliminating the need for a year-long wait. To improve upon our research model and better mimic human conditions, we plan to extend the testing duration by incorporating additional time points up to at least one year in future studies. Furthermore, the prolonged effects of SARS-CoV-2 on mental health necessitate further investigation.

Second, we used a murine model, and findings in rodents may not fully translate to human conditions. Future research should explore the temporal dynamics of microglial activation and neuroinflammation throughout different stages of SARS-CoV-2 infection.

Third, we identified that brain infection in mice developed mild respiratory diseases. We did not further evaluate the case of asymptomatic infection because a clinical study suggested that the risk of developing long-term symptoms in asymptomatic SARS-CoV-2 infected persons was significantly lower than those in symptomatic SARS-CoV-2 infection cases (70).

Fourth, we utilized a mouse-adapted SARS-CoV-2 strain that shares some key spike mutations with the Beta variant. However, the specific potential, impact, and severity of neuroinvasion resulting from different SARS-CoV-2 variants, such as the Omicron variant, and the potential effects of breakthrough infections in vaccinated individuals, remain areas that require further investigation to provide a comprehensive understanding.

Fifth, we did not systematically investigate the potential impact of elevated cytokines and chemokines induced by respiratory COVID-19 on microglial activation and their potential effects on impairing CNS neuron function. It is well-established that exposure to chemotherapy drugs, brain radiation, or systemic inflammation can lead to persistent activation of certain microglia (1, 71–74). Interestingly, a recent study of SARS-CoV-2 infection in AAV-hACE2 sensitized mice or H1N1 influenza infection in mice revealed that mild respiratory disease-activated white-matter-selective microglia, leading to oligodendrocyte loss, impaired neurogenesis, and elevated CCL11 levels (75). These findings highlight the potential impact of systemic inflammation on microglia activation and brain function. Longitudinal studies in human cohorts are necessary to establish causality between viral neurotropism, microglial activation, and neuropsychiatric outcomes. Moreover, investigating potential sex-dependent differences in behavioral responses and microglial activation can provide a more comprehensive understanding of COVID-19-associated neuropsychiatric sequelae. Addressing these limitations through further research is essential to advance our understanding of the complex neurological effects of SARS-CoV-2 infection and its variants. By exploring these areas further, we can gain valuable insights to develop more effective interventions and treatment strategies for individuals affected by COVID-19 and its neurological consequences. The evaluation of the consequences of N501Y_MA30_ infection on the CNS has provided valuable insights, positioning this mouse-adapted strain as an auspicious model for studying the neurological manifestations of PASC. Continuation of research utilizing this mouse model, alongside other relevant models, is pivotal in advancing our understanding of the neurological effects of SARS-CoV-2 and facilitating the development of effective interventions for PASC.

Finally, we utilized bulk RNA-sequencing to analyze gene changes following viral infection. However, this approach may not fully capture gene changes in specific cell types, such as microglia, which could potentially confound the observed neuron-microglia interactions. To precisely elucidate the immune response and subsequent changes in neuronal activity genes, future studies should consider supplementing with single-cell RNA-seq focusing on specific brain regions.

## Conclusion

Our study demonstrates that SARS-CoV-2 neurotropism induces anxiety- and depression-like behaviors through the activation of microglia and subsequent neuroinflammation. These findings underscore the importance of considering mental health disturbances in COVID-19 patients and emphasize the need for integrated approaches to address both the physical and mental health aspects of the pandemic. Targeting microglial activation and neuroinflammatory pathways may offer promising therapeutic avenues to mitigate the neurobehavioral consequences of SARS-CoV-2 infection. Further investigations in human cohorts are warranted to validate and extend our preclinical findings for potential translational applications.

## AUTHOR CONTRIBUTIONS

J.D. and K.L. conceived the project. J.D., K.L., and Q.G. designed the experiments. Q.G. and J.D. performed the patch-clamp and immunostaining experiments and data analysis. S.Z. and K.L. performed the behavior and molecular experiments and data analysis. K.L., J.D., and Q.G. wrote the manuscript. All authors reviewed and edited the manuscript.

## ACKNOWLEDGMENTS

J.D. is supported by the National Institutes of Mental Health (1R01MH113986), the Cystic Fibrosis Foundation (002544I221), and the University of Tennessee Health Science Center start-up fund. K.L. is supported by the Cystic Fibrosis Foundation and the Cleveland Clinic Florida Research and Innovation Center start-up fund.

---

**Supplemental Figure 1.**
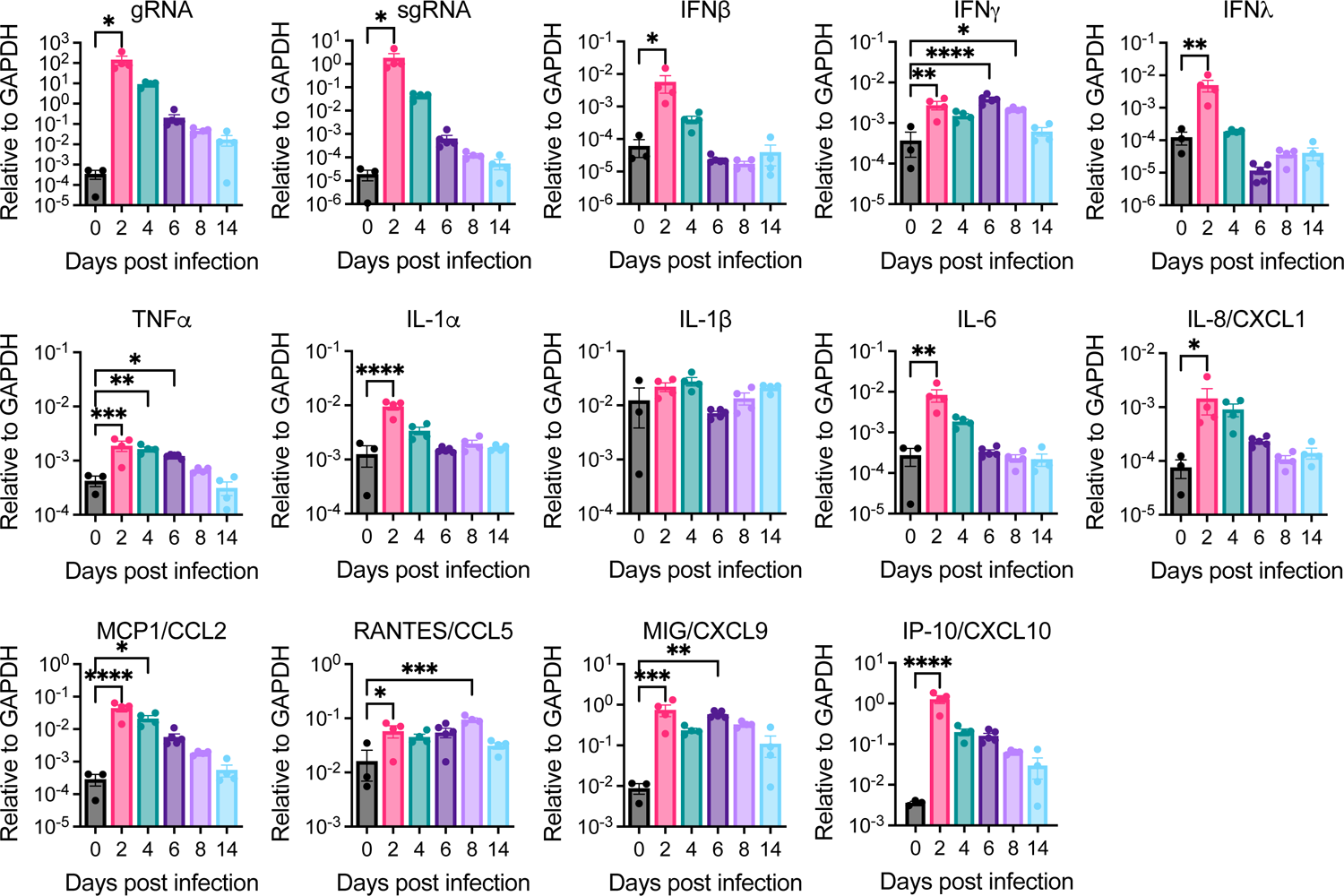
SARS2-N501Y_MA30_ Virus RNA Replication and Host Immune Responses in the Lungs. Total RNA was extracted from lungs collected from C57BL/6 mice (n=3 to 5), intranasally infected with 10^4^ PFU of SARS2-N501Y_MA30_ at designated days post-infection. Viral RNA, as well as host cytokine and chemokine transcripts, were quantified using quantitative real-time polymerase chain reaction (qRT-PCR). Viral gRNA or sgRNA CT values from mock-infected tissues (0 dpi) exceeded 35. Transcript levels were normalized to GAPDH and presented as 2^-ΔCT. Statistical significance was assessed using ordinary one-way ANOVA, with significance levels indicated as follows: *p < 0.05, **p < 0.01, ***p < 0.001, ****p < 0.0001. Data are presented as mean ± SEM.

**Supplemental Figure 2.**
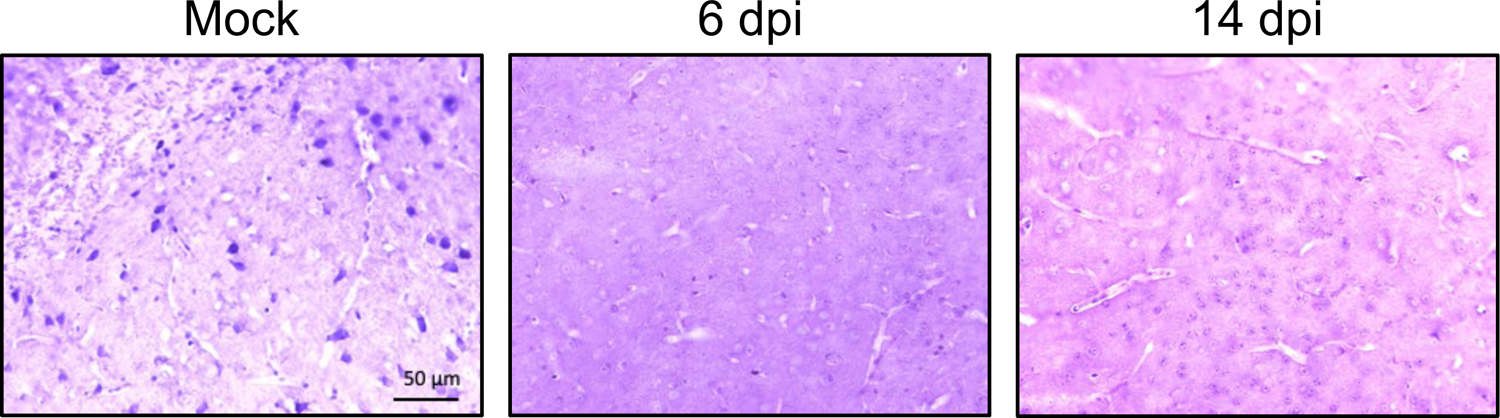
Brain Histopathology after Mock and SARS2-N501Y_MA30_ Infection. The representative images of brain tissue sections were stained with hematoxylin and eosin (H&E) at indicated days following either mock or SARS-CoV-2 infection. Scale bar: 50 μm

**Supplemental Figure 3.**
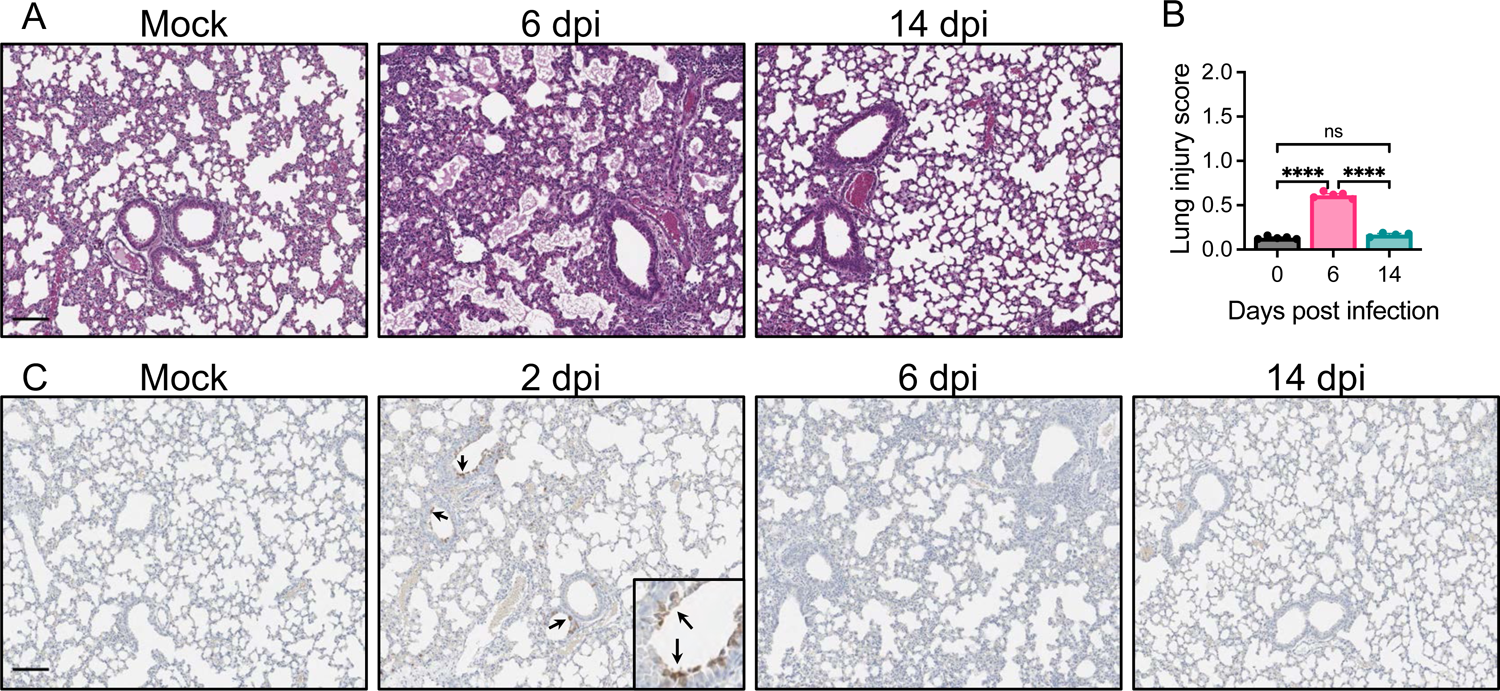
Lung Pathology and Immunohistochemistry of Viral Protein. **(A)** Lung sections from mice infected with SARS2-N501Y_MA30_ (n=5 for mock infection and for 6 dpi; n=4 for 14 dpi) underwent H&E staining. **(B)** Acute lung injury severity was assessed blindly following ATS guidelines (32), with 0 for no injury, 1 for mild to moderate injury, and 2 for severe injury. ****p < 0.0001. Scale bars: 100 µm. Data are shown as mean ± SEM. **(C)** Lung Tissue Immunohistochemistry revealing SARS-CoV-2 N protein (black arrows). Representative images show the presence of SARS-CoV-2 N protein (indicated by black arrows) at specified days post-infection. Scale bar: 100 μm.

**Supplemental Figure 4.**
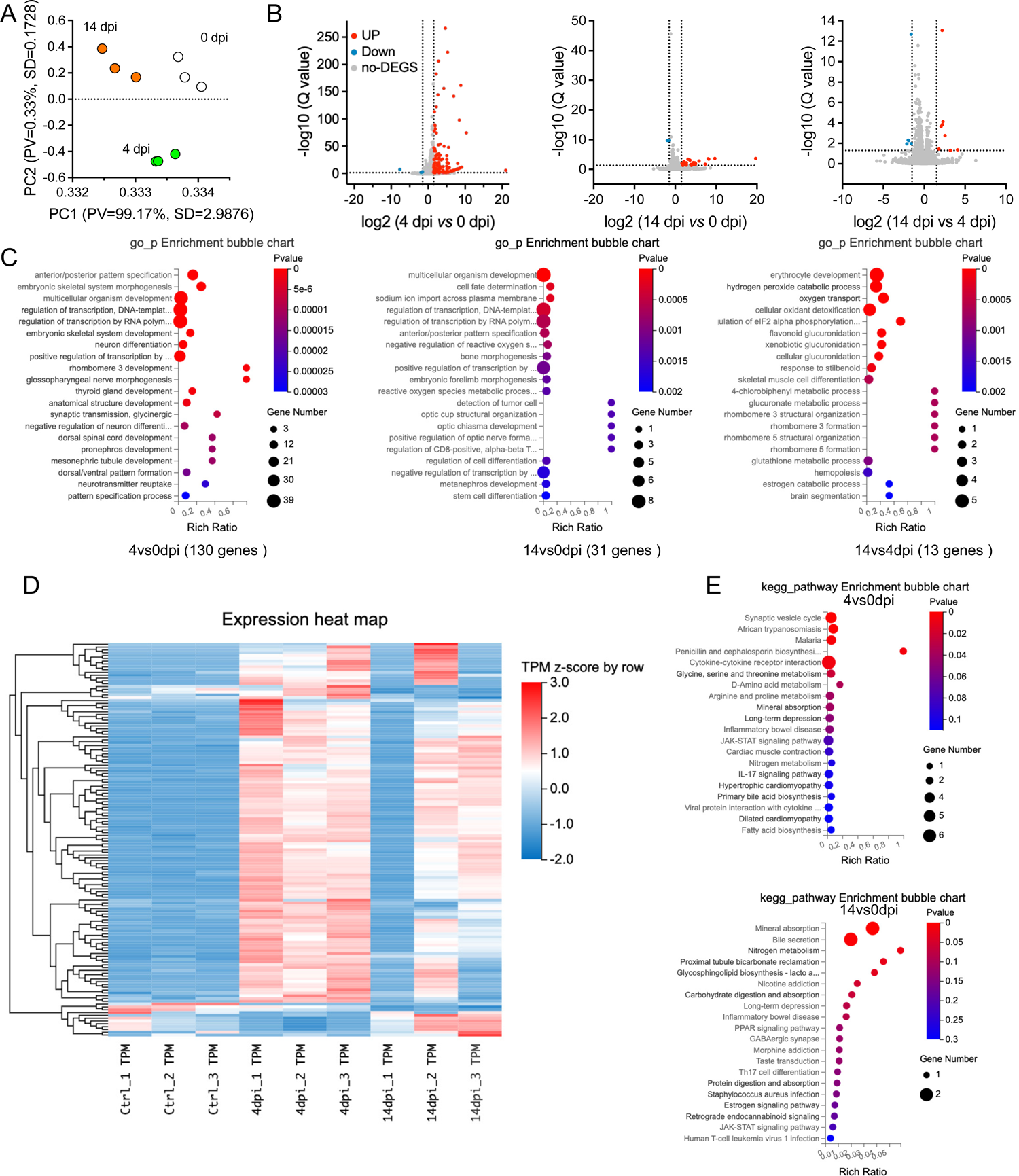
Transcriptomic Profile Overview in SARS2-N501YMA30 infected mouse brains. **(A)** Principal Component Analysis (PCA) of transcriptome profiles comparing mock-infected brains (0dpi) with those at 4 and 14 days post-infection. **(B)** Volcano Plot Analysis: The x-axis denotes log2-transformed fold changes, and y-axis indicates -log10-transformed significant values. (Thresholds set at |log2fold| ≥ 1.5, Q value ≤ 0.05) **(C)** Gene Ontology (GO) Enrichment Analysis of DEGs across groups. **(D)** Heatmap visualization of 140 DEGs identified post-infection. **(E)** Kyoto Encyclopedia of Genes and Genomes (KEGG) Pathway Analysis comparing groups at 4dpi vs 0dpi, 14dpi vs 0dpi.

**S1 Table.**
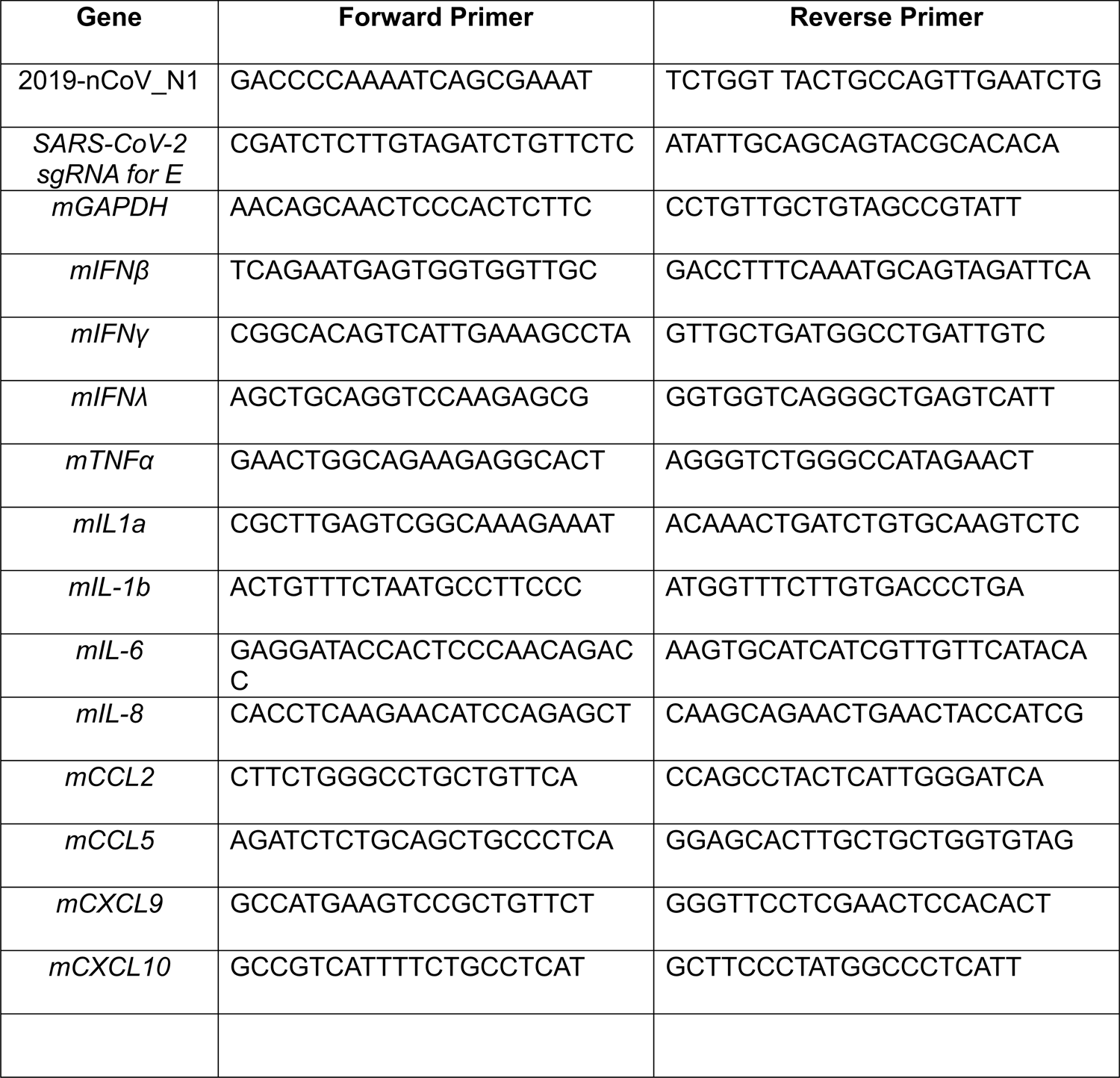

